# Genomic dialects: How amino acid properties and the second codon base shape the informational accents of life

**DOI:** 10.64898/2026.04.21.720023

**Authors:** Octavio Martínez, Neftalí Ochoa-Alejo

## Abstract

Codon Usage Bias (CUB) is a fundamental feature of genomic architecture, reflecting a balance between mutational pressure and natural selection. We propose a “genomic dialects” framework, where species-specific CUB profiles represent “informational accents” constrained by biochemical and structural requirements. Utilizing a normalized informational index based on Shannon’s entropy, we analyzed CUB profiles for 18 amino acids across 1,406 species from the three domains of life. Linear models were employed to investigate the relationship between CUB and physicochemical properties, including Saier’s second-codon-base classification, molecular volume, hydrophobicity, aliphatic/aromatic status, and dissociation constants. CUB distributions are highly skewed, with *>* 52% of values below 0.1, suggesting a near-optimal use of the genetic code’s potential. We demonstrate that amino acid properties significantly influence CUB, with Saier’s classification explaining up to 69% of variance in Archaea and *≈* 47% across all taxa. Hydrophobic amino acids (*Q*_1_ class) consistently exhibit higher average CUB than hydrophilic ones, particularly in microbes. Individual species models reveal extreme correlations; for example, in the alga *Chlamydomonas reinhardtii*, Saier classes explain *>* 95% of CUB variance. Finally, we show that CUB-based dendrograms represent phenetic similarity (“genomic accents”) rather than reliable phylogenetic reconstructions, as they rarely coincide with the true Tree of Life. Our findings indicate that the “rules” of genomic dialects are largely anchored in the dual requirements of translational fidelity and protein stability. The observed “informational accents” are proximately governed by the metabolic and genomic machinery under the constraints of the drift-barrier hypothesis. This study provides a robust framework for understanding how the physical realities of amino acids have shaped the evolution of the genetic code’s informational use across the tree of life.

## Introduction

Codon Usage Bias (CUB) is measured on a numeric scale from zero to one. A value of zero indicates that all synonymous codons for a given amino acid (*aa*) appear with equal frequency within the CDSs of a genome, whereas a value of one is reached when only a single codon is used exclusively for that *aa*. Consequently, for any given species, we obtain a vector of numeric CUB values for the 18 *aa*—including the ”Stop” signal—that possess more than one codon in the genetic code. This CUB profile summarizes the species-specific patterns of codon usage, highlighting the unique particularities in how that organism utilizes the genetic code (See also **S1.1** in S2 Text).

The systematic variation in CUB across taxa suggests that the genetic code is not merely a static lookup table, but rather a dynamic system of “genomic dialects”. This “accent” metaphor aligns with the Cell Language Theory proposed by [1], which suggests a formal isomorphism between the rules of linguistics and molecular biology. Just as human accents are defined by specific phonotactic constraints-rules that govern which sounds are preferred or permitted in a language, different taxa exhibit specific “codon-tactic” preferences.

This view is supported by the work of [2], who noted that both systems rely on a hierarchy of discrete units to convey complex information. In this context, CUB represents the “organic semantics” of a species [3], where the choice of a synonymous codon is not neutral, but an expression of the organism’s unique informational and biochemical environment.

### CUB measurement

The estimation of CUB has a long history, beginning with early analyses of bacterial genomes [4, 5]. While the Relative Synonymous Codon Usage (RSCU) index [6] remains useful for assessing selection in highly expressed genes, a universal framework requires a measure that is independent of reference sets or gene expression levels.

Wright’s “Effective Number of Codons” (ENC) index [7] provides such a measure, typically ranging from 20 to 61. To facilitate a more intuitive comparison across different amino acids and taxa, we utilized a CUB index, *B_ik_*, that is normalized to vary from 0 to 1:

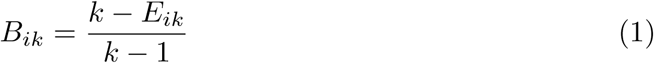

where *k* represents the number of synonymous codons and *E_ik_*= 2*^Hik^* is the effective number of codons derived from Shannon’s entropy (*H*) and was implemented in our R package “Shannon.codon” [8]. For example, in humans, the *aa* Gln is encoded by two codons with observed frequencies of **p** *≈* (0.27, 0.73). This distribution yields *E_Gln_ ≈* 1.79, resulting in a calculated bias of *B_Gln_ ≈* 0.21 (See also **S1.3** in S2 Text).

The mathematical properties and the biological relevance of the *B_ik_*index were previously established and validated in [9]. In that study, we demonstrated that this normalized informational measure effectively captures the nuances of codon usage across a broad taxonomic range, providing a robust basis for comparing “informational accents” throughout the tree of life. The current work extends this framework by studying with detail the implications of the differences in CUB measures between the three life domains to further explore the biochemical and structural drivers of these genomic dialects.

### The biochemical determinants of genomic dialects

While the existence of CUB is well-documented, the factors that shape these “informational accents” across diverse taxa remain a subject of active debate. If the genetic code is indeed a language, its rules must be constrained by the physical and chemical realities of the molecules it encodes. In this study, we hypothesize that the variance observed in CUB is not fully stochastic but is fundamentally anchored in the biochemical properties of amino acids, as for example their hydrophobicity and molecular volume.

In [10], the author made the insightful observation that the second position in the codon (denoted here as *P* 2), is strongly correlated with physiochemical *aa* types, likely reflecting a primordial code that partitioned residues into hydrophobic, hydrophilic, and semipolar blocks to minimize the impact of early translational errors.

The Saier classification provides a robust framework for understanding these genomic dialects by highlighting several deterministic rules that govern the code’s architecture. Beyond the primary influence of *P* 2, the code employs a strategy of error-minimization where amino acids with similar properties are assigned to codons differing by only a single nucleotide, effectively reducing the deleterious impact of mutations on protein stability. This stability is further reinforced by the asymmetric hydrogen-bond strength between mRNA and tRNA, where pyrimidines in the mRNA (such as Uracil or Cytosine) form more stable pairings than when the corresponding purine is present. Furthermore, the importance of the third “wobble” position (*P* 3) is not random but is strictly dictated by the identity and bonding strength of the nucleotides at *P* 1 and *P* 2. Finally, the selection of rare synonymous codons serves a functional “phonotactic” purpose by programming translational pauses that facilitate proper co-translational protein folding.

Building upon these principles, here we expand and explain in detail the informational properties first discussed in [9], with a specific emphasis on how these phonotactic constraints manifest in CUB values across the tree of life. By analyzing these properties through the lens of the Saier classification, we aim to demonstrate that the rules of a genome are largely determined by the dual requirements of translational fidelity and protein stability.

## Materials and methods

Table 1 presents the main groups of organisms studied here.

**Table 1.** Groups of species included in this study.

| Domain | Kingdom | Key | Number |
| --- | --- | --- | --- |
| Archaea |  | Ar | 435 |
| Bacteria |  | Eb | 126 |
|  | Animalia | An | 652 |
| Eukarya | Plantae | Pl | 122 |
|  | Fungi | Fu | 71 |

The total number of species included in this study is 1,406, of which 845 are eukaryotes. Each group is identified by its “**Key**”, which is consistently used throughout the figures and tables. Within each group, individual species are uniquely labeled by their respective key followed by a sequential number. The full scientific name for each species is available in the “Shannon.codon” R package [8] as well as in the Supporting information S3 R.

In Table 1 we included representative species from the three life domains, 435 Archaea, 126 Bacteria –represented only by species from the Enterobacteriaceae family of Gram-negative bacteria, and 845 eukaryotes, which are sub-classified into the three main kingdoms; 652 animals, 122 plants and 71 fungi. We exclude from the groups in Table 1 21 viruses –which do not have a shared evolutionary ancestor like cellular life, four protists (amoebas) –which are traditionally considered as a fourth kingdom but for which we do not have a representative sample size, and three bacteria which do not belong to the Enterobacteriaceae family.

Codon frequency data were downloaded from the Codon Statistics Database [11], as described in [9] and are available as supporting information **S1** on that paper. CUB values for the 18 *aa* in all species were obtained by equation (1) using our R package “Shannon.codon” [8] and are included here as S1 Dataset.

To detect the effect of *aa* characteristics on CUB values, we fitted linear models to individual or grouped CUB values as a function of *aa* properties. The variables considered included the Saier classification (*P* 2 quadrant), the number of encoding codons (*n.cod*), van der Waals radii (*vWvol*), hydrophobicity (*H_y_*), aliphatic/aromatic classes (*A_c_*), and dissociation constants (*pK*_1_ and *pK*_2_).

Table 2 presents the values of the variables for each one of the 18 *aa* which are coded by more than one codon (See also **S1.2** in S2 Text).

**Table 2.** Values of variables per amino acid (*aa*).

| $aa$ | $P2$ | $n.cod$ | $vWvol$ | $H_y$ | $A_c$ | $pK_1$ | $pK_2$ |
| --- | --- | --- | --- | --- | --- | --- | --- |
| Ile | $Q_1$ | 3 | 124 | Pho | Al | 2.32 | 9.76 |
| Leu | $Q_1$ | 6 | 124 | Pho | Al | 2.33 | 9.74 |
| Phe | $Q_1$ | 2 | 135 | Pho | Ar | 2.20 | 9.31 |
| Val | $Q_1$ | 4 | 105 | Pho | Al | 2.39 | 9.74 |
| Ala | $Q_2$ | 4 | 67 | | Al | 2.35 | 9.87 |
| Pro | $Q_2$ | 4 | 90 | | | 1.95 | 10.64 |
| Ser | $Q_2$ | 6 | 73 | | | 2.19 | 9.21 |
| Thr | $Q_2$ | 4 | 93 | | | 2.09 | 9.10 |
| Asn | $Q_3$ | 2 | 96 | Phi | | 2.14 | 8.72 |
| Asp | $Q_3$ | 2 | 91 | Phi | | 1.99 | 9.90 |
| Gln | $Q_3$ | 2 | 114 | Phi | | 2.17 | 9.13 |
| Glu | $Q_3$ | 2 | 109 | Phi | | 2.10 | 9.47 |
| His | $Q_3$ | 2 | 118 | Phi | Ar | 1.80 | 9.33 |
| Lys | $Q_3$ | 2 | 135 | Phi | | 2.16 | 9.06 |
| Tyr | $Q_3$ | 2 | 141 | Phi | Ar | 2.20 | 9.21 |
| Arg | $Q_4$ | 6 | 148 | | | 1.82 | 8.99 |
| Cys | $Q_4$ | 2 | 86 | | | 1.92 | 10.70 |
| Gly | $Q_4$ | 4 | 48 | | | 2.35 | 9.78 |

$P2$ quadrant: $Q_1$ (U) Hydrophobic, $Q_2$ (C) Semi-polar, $Q_3$ (A) Hydrophilic, $Q_4$ (G) Semi-polar+Arg.
Hydrophobicity, $H_y$ : “Phi” in Hydrophilic $aa$ , “Pho” in the Hydrophobic ones.
Aliphatic / Aromatic class, $A_c$ : “Al” for aliphatic $aa$ , “Ar” for aromatic ones.
Dissociation constants: $pK_1$ for the carboxyl ( $-\text{COOH}$ ) terminus while $pK_2$ for the amino ( $-\text{NH}_2$ ) terminus.
Source for $P2$ : [10] and for $vWvol$ , $H_y$ , $A_c$ , $pK_1$ and $pK_2$ : [12].

In Table 2 we can see that there are qualitative (*P* 2, *H_y_* and *A_c_*) as well as quantitative (*n.cod*, *vWvol*, *pK*_1_ and *pK*_2_) variables for the 18 *aa* encoded my more than one codon in the nuclear genetic code. Variable *P* 2 gives the Saier classes (see **FIG 3** in [10]), which label the 18 *aa* into 4 categories; 4 in *Q*_1_ which have the U base in the second position and are hydrophobic, 4 in *Q*_2_ which have the C base in the second position and are semi-polar, 7 in *Q*_3_ which have the A base in the second position and are hydrophilic and, finally, 3 in *Q*_4_ which have the G base in the second position and are also classified as semi-polar but include also the *aa* Arg as a kind of “outlier” in that class. The merit of the Saier classes, denoted as variable *P* 2 in Table 2, is that it links a genetic variable (the second position in the codons), with very relevant biochemical *aa* characteristics, allowing the study of CUB values as function of those classes.

**Fig 1.**
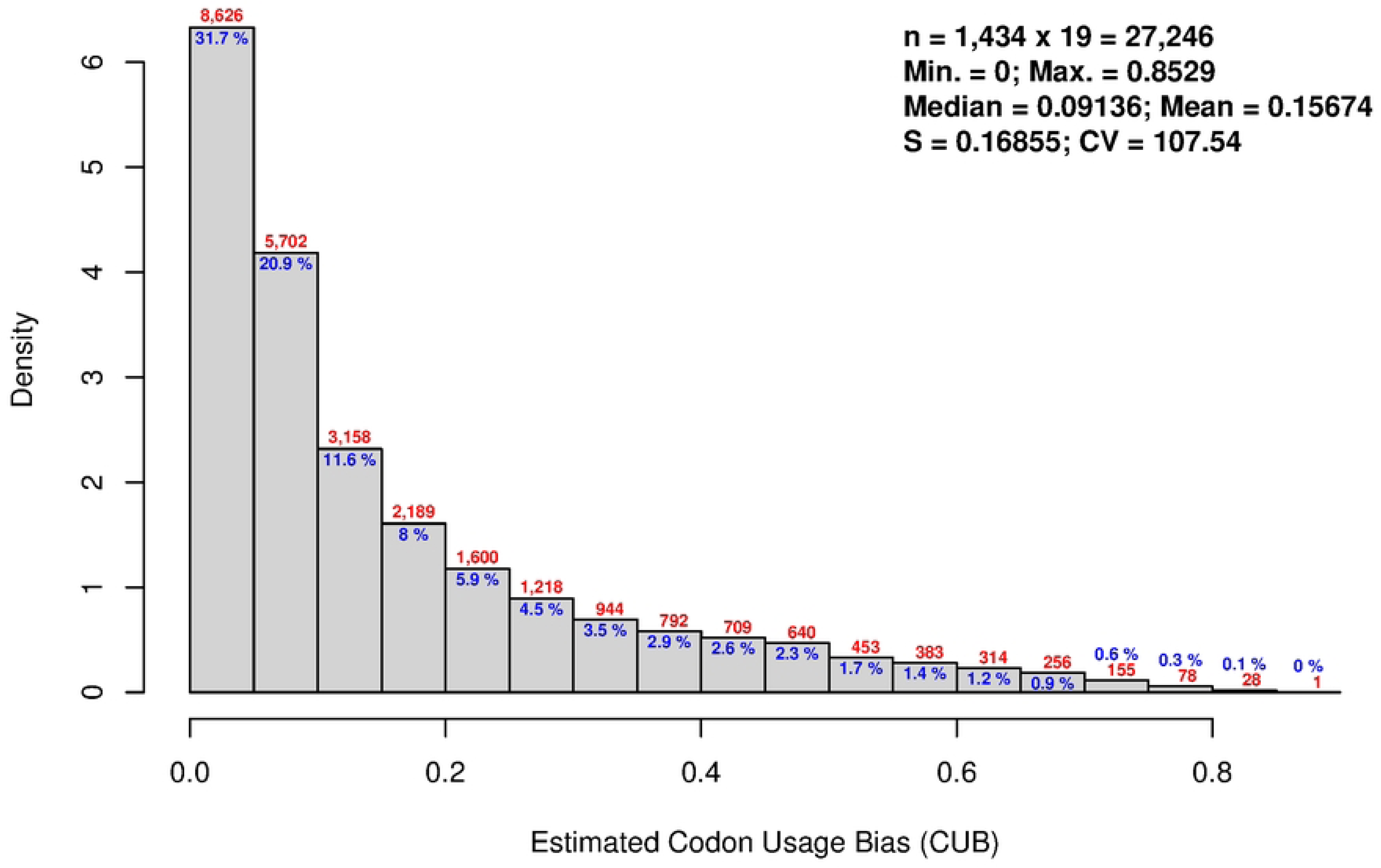
Histogram for the distribution of CUB values. Figures in red and blue give the number and percentages of values per bar, respectively. In legend “S” is the standard deviation and “CV” the Coefficient of Variation; CV = 100 *×* (S/mean).

**Fig 2.**
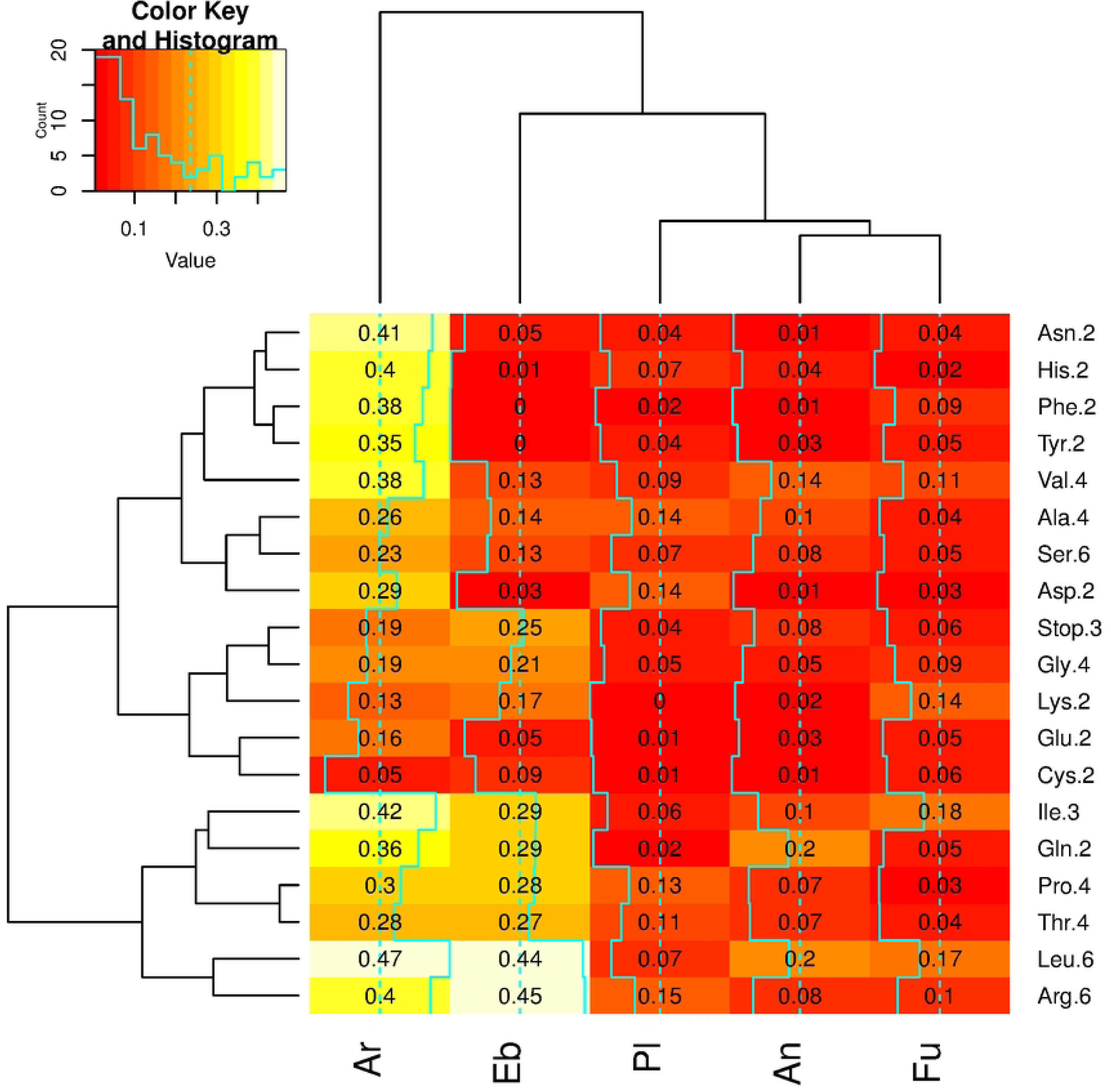
Heat map for the median CUB values per main group and *aa*. *X*-axis presents the main groups of species while *Y* -axis have the *aa*. *aa* abbreviations are followed (after the full stop) by the number of codons which code for each *aa*.

**Fig 3.**
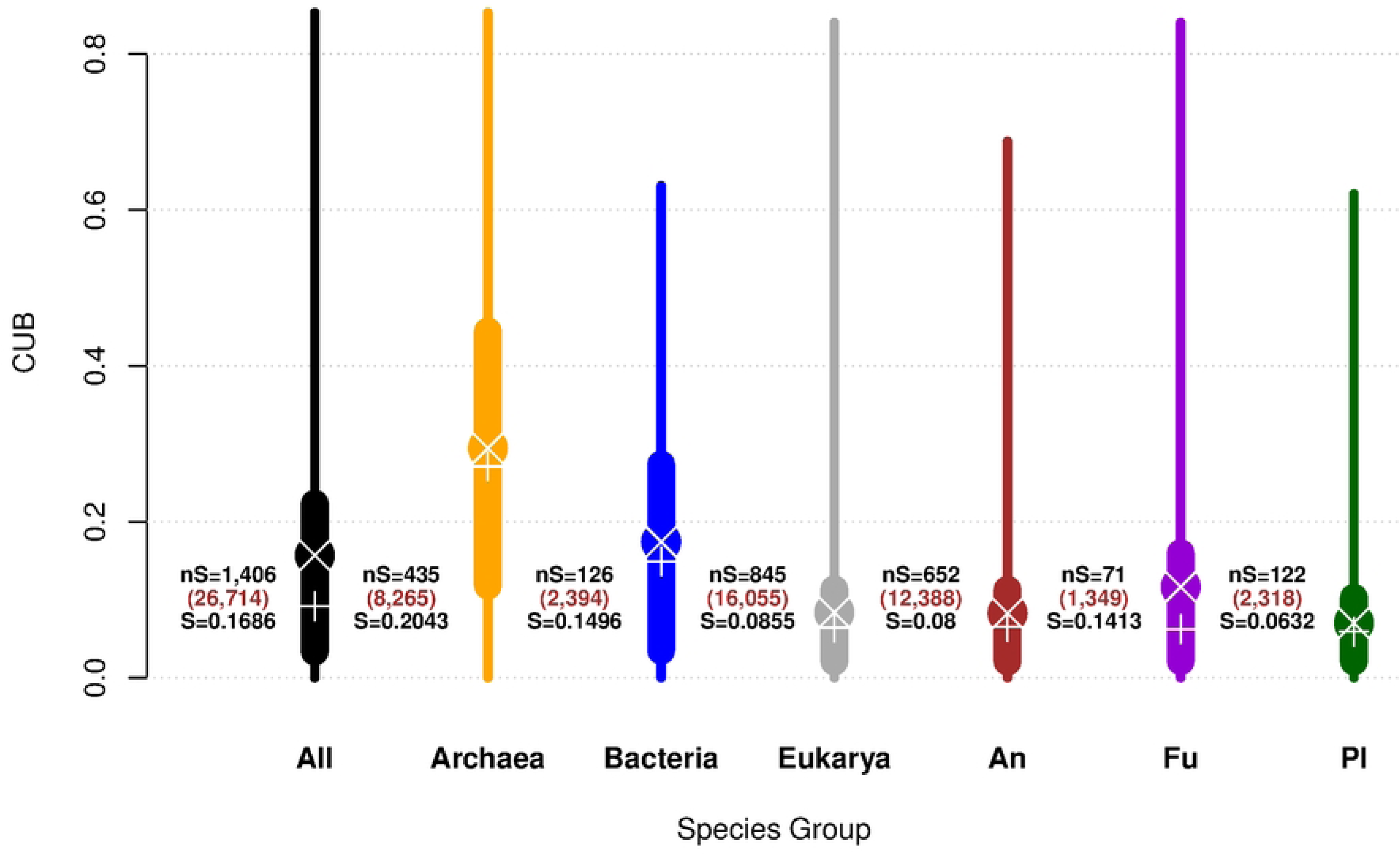
Statistics for CUB by group. Narrow lines for each group go from the minimum of the CUB value (0 in all 7 cases) up to the maximum per group, while wide lines per group contain the interquartile range, i.e., values between the quantiles Q 0.25 and Q 0.75. Means and medians per group are shown by white crosses and plus signs, respectively. Annotations per group at the left are number of species, nS, number of CUB values are between parenthesis and standard deviations, per group, *S*.

Variable *n.cod* in Table 2 takes values 2, 3, 4 and 6 for 9, 1, 5 and 3 *aa*, respectively –but let out the 3 codons, TAA, TAG and TGA, that encode the “Stop” signal. We can also investigate if there is a relation between the quantitative variable *n.cod* and the values that CUB takes in different species.

On the other hand, variable *vWvol* in Table 2, the van der Waals radii of amino acids, give us the effective *aa* size, which is crucial for analyzing protein structure, packing, and surface area. By modeling CUB values as function of the *aa* size (*vWvol*) we can investigate if those variables are significantly related in each one of the species groups.

Variable *H_y_* in in Table 2 recodes hydrophobicity, but exclusively for 11 of the 18 *aa*, classifying 4 of them as hydrophobic (“Pho”) and 7 as hydrophilic (“Phi”), but letting out of this classification the remaining 7 *aa*. Models of CUB as function of *H_y_* will then consider only 11 values per species, isolating exclusively the hydrophobicity effect on CUB.

On the same lines, variable *A_c_* has two categorical values, “Al” for the 4 aliphatic *aa* and “Ar” for the 3 aromatic ones; i.e., only 7 *aa* are taken into account in the *A_c_*, allowing the modeling of CUB by taking into account only the aliphatic or aromatic nature of the *aa*.

Finally, the dissociation constants, *pK*_1_ –for the C-terminus, and *pK*_2_ –for the N-terminus of each *aa*, give numeric variables which are worth exploring to investigate if they present a relation with the CUB values of the *aa* in different species.

Different model parameterizations were evaluated to explore the effects of the variables presented in Table 2 over CUB values. This was performed across various settings, including models segregated by taxonomic group (Table 1) and aggregate models excluding taxonomic affiliation.

Additionally, to explore the taxonomic consistency of dendrograms constructed from CUB values, we employed the R package “DendroLikeness” [13]. This allowed us to compare the frequencies of dendrograms obtained from random species samples against the topology expected from the known Tree of Life (Table 1).

All calculations were performed in R [14] version 4.3.2 on the aarch64-apple-darwin20 (64-bit) platform. Detailed descriptions of the analyses are provided in S2 Text, and the relevant R objects are publicly available (see S3 R).

## Results and Discussion

### Distribution of estimated CUB values

CUB values measure in an inverse way the informational optimality in the use of the genetic code, because CUB values close to zero imply the best use of the coding potential –all synonymous codons are employed in approximately the same frequency, while values close to one imply that the coding potencial is “wasted” –because only one of the possible codons is used. Given that we have 18 *aa* coded by more than one codon (Table 2), plus the 3 codons that encode the “Stop” signal, thus each one of the species studied yields a total of 19 CUB values to be considered. Fig 1 shows the histogram of all estimated CUB values.

In Fig 1 we can see that the rank of estimated CUB values includes 90% of the possible values of the CUB coefficient, which goes from zero to one –equation (1). However, the CUB distribution is extremely skewed, with most data points clustered on the left hand side. In fact, more than half of the CUB values are smaller than 0.1 (percentages in the first two left side bars; 31.7 + 20.9 = 52.6%), demonstrating that life forms tend to make a near optimum use of the genetic code potential.

The evolutionary forces that shape the CUB profile in a species are restricted in principle by the proteome, the set of proteins that must be expressed by the organism to survive. Within these evolutionary forces, mutation is the main random component, followed by the effective population size which vary due to environmental factors. If the CUB distribution shown in Fig 1 is representative of all life diversity, we could conclude that the genetic code has a near optimum informational use of a “language” that was definitely fixed at the dawn of life.

Now we proceed to study the “accents” of the use of the genetic code, which are reflected in the differences of CUB profiles per species’ group. Fig 2 presents a heat map estimated from the median CUB values per species group and *aa*.

Dendrograms on the margins were constructed by the complete method from the medians of CUB values per group of species and *aa*. Figures in the cells give the corresponding CUB median.

In Fig 2 we used the median instead the mean of CUB values to avoid the strong influence of CUB outliers, however **S2.1** in S2 Text presents and discuss box plots and heat maps using different measures.

Currently, the true topology of the phylogenetic relations between the groups studied here is *T_t_* = *{*Eb, *{*Ar, *{*Pl, *{*Fu, An*}}}}* [15], in which eukaryotes (Pl, Fu and An) are within archaea, and there is a close relationship between animals and fungi, previously inferred in [16]. In contrast, the topology observed in the upper margin in Fig 2 is *T_o_* =*{*Ar, *{*Eb, *{*Pl, *{*An, Fu*}}}}*. Both topologies, *T_t_* and *T_o_*, coincide if forming an inner group with the eukaryotes, “*{*Pl, *{*Fu, An*}}*”, and in both cases animal and fungi are clustered together first and then plants joins that group. This fact indicates that within eukaryotic organisms the median of CUB values is a reasonable proxy for phylogenetic relations. However the true topology, *T_t_*, implies that bacteria –represented here by Eb, is the group basal to the tree of life, while in Fig 2 the observed topology, *T_o_*, set archaea (Ar) as the basal group. This results from the fact that median CUB values in Eb are closer to the ones for eukaryotes than to the ones for archaea, and that can also be noticed by the change in color of the heat map which, with few exceptions, shifts from yellow (large values) on the left for Ar to more reddish tones on the right (smaller values).

The left hand side dendrogram in Fig 2 clusters the *aa* by their likeness in median CUB values. In that dendrogram the two *aa* with more alike median CUB values are Pro and Thr, both of them encoded by two codons in the genetic code. Nevertheless, there is not a high congruence with the likeness between *aa* and the number of codons coding for them; in fact the dendrogram in Fig 2 has two well segregated clusters which show an heterogeneous number of codons. The group in the lower left hand side of that dendrogram is formed by the *aa* Ile, Gln, Pro, Thr, Leu and Arg, which includes the two *aa* coded by 6 codons, Leu and Arg, two of the four coded by two codons, Pro and Thr, but also one coded by 2 codons, Gln and one coded by 3 codons, Ile.

A more detailed perspective of the differences of CUB values can be obtained by examining Fig 3, which presents the statistics for CUB by group, including all taxonomic divisions in Table 1.

The first plot in Fig 3, in black for “All”, is a different way to present the distribution of CUB values, previously seen as an histogram in Fig 1 (which also included CUB values out of the main groups). In the second plot in Fig 3, in orange for “Archaea”, we can appreciate how this life domain presents the highest mean and median CUB values, being also the most variable group, with a CUB standard deviation *S ≈* 0.2, which is also the largest of the divisions presented in Fig 3. This is consistent with the results presented for Archaea in [9], which showed that this branch of the tree of life is the most diverse on informational codon use.

On the other hand, the group which presents smaller values of CUB central tendency as well as smaller variation is plants (Pl), the last plot in Fig 3 in green at the right hand side. This suggests that during the evolution of this clade CUB values have been more conserved than in other groups. See also **S2.2** in S2 Text.

### Correlating CUB values with aa characteristics

We have demonstrated how the universal genetic code is used with different “*accents*” (CUB profiles) by the various groups of organisms. We are now in the position to ask if there are *aa* characteristics associated with CUB values.

When discussing putative answers to that question we must keep in mind the fact that correlation does not imply causation; i.e., we must refrain from assigning causality to changes on average CUB values to a given *aa* characteristic, even if the differences found are very large and also with a very high statistic significance (*p*-values *≈* 0). To consider that an *aa* attribute *causes* a change in average CUB value we must have not only robust statistical evidence, but also a strong chain of molecular arguments that will explain such change. Given that the exact history of CUB changes along any lineage is unknown, it appears impossible to segregate random from causal factors on CUB values. Nevertheless, the study of the relative sizes of of CUB changes given by different *aa* characteristics could give some insights on the modification in the use of the genetic code which occurred during the evolutionary history of the different species’ groups.

### Average CUB values by group and Saier classes

In Table 2 we have seen how the 18 *aa* are classified into 4 classes, defined by the second position of the codon, *P* 2 = *{Q*_1_*, Q*_2_*, Q*_3_*, Q*_4_*}*, and presented in [10]. To analyze the effects of the Saier’s class on the average CUB value we fitted linear models CUB = *β_i_Q_i_*, for each one of the groups presented in Table 1 –including the set of all species, and then standardized the *β* coefficients to be able to compare their values in a uniform scale. Fig 4 presents a graphic summary of the results for the ANOVA’s of the models.

**Fig 4.**
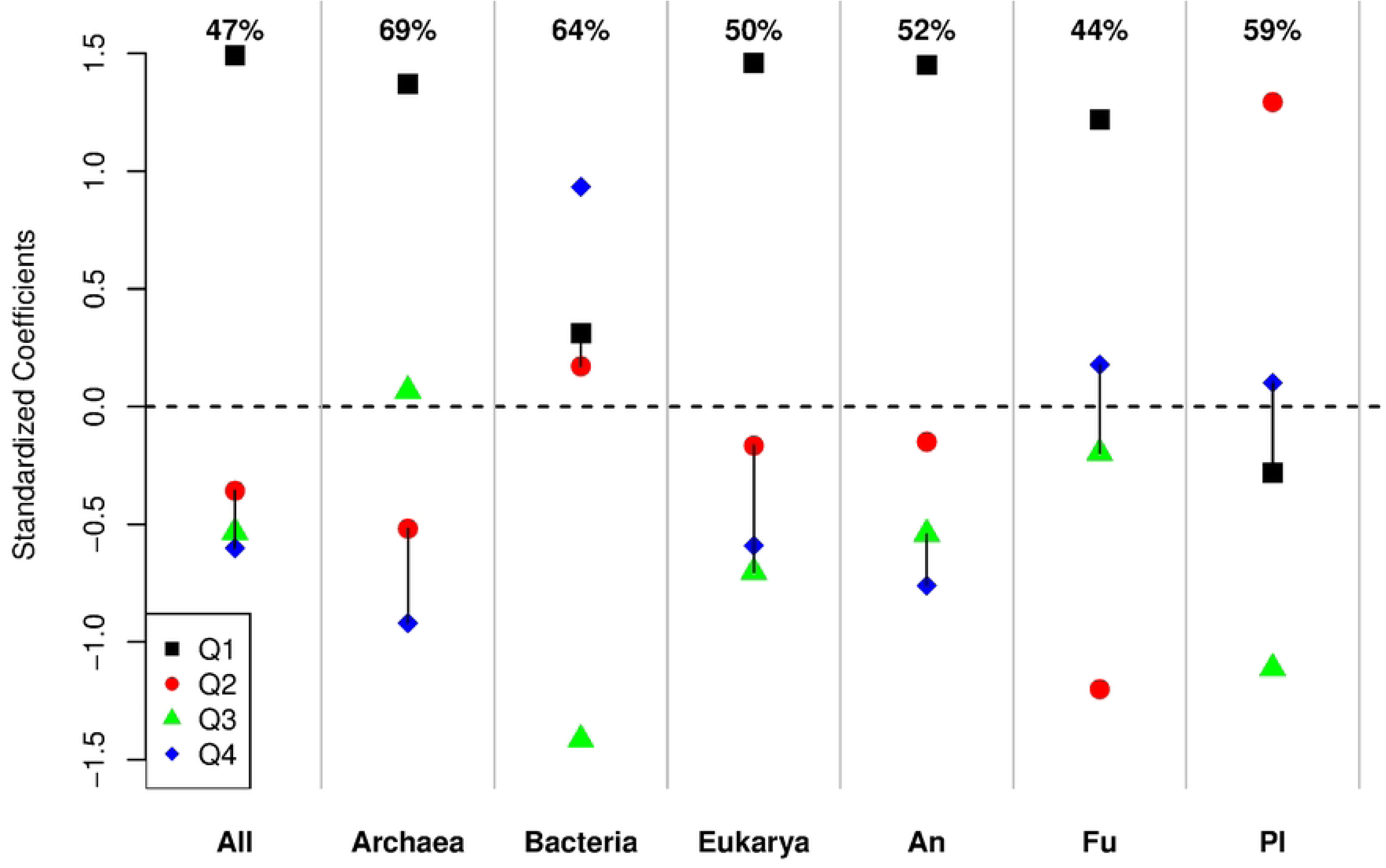
ANOVA results for CUB by Saier classification. Plot of standardized coefficients (*Y* -axis) for the ANOVA of the models “CUB = *β_i_Q_i_*” by group of species (*X*-axis). Numbers at the top give the percentages of CUB variance explained by each model, %*V* = 100 *×* 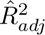. Not significant differences in Tukey tests (adjusted *p*-values *>* 0.01) are linked by a vertical line.

In Fig 4 the *Y* -axis gives the standardized values of the *β* coefficients for the models CUB = *β_i_Q_i_* using different symbol and color for each *Q_i_* (see legend). Given the parametrization of the models, the estimated ̂*β_i_*’s give the averages CUB values for the corresponding classes, *Q_i_*’s. Results of estimates per species group are shown along the *X*-axis, and the sum of standardized ̂*β_i_*’s coefficients per group of species adds to one in every column, allowing the comparison of the models in the same scale. Additionally, the results of Tukey’s tests are shown by linking with a vertical lines the symbols for the *Q_i_*’s that were not significantly different using an adjusted *p* value of 0.01 while the figures at the top of each line show the percentage of CUB variance explained by the corresponding model, %*V* = 100 *×* 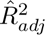, where 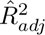 is the adjusted determination coefficient of the corresponding model.

In Fig 4 we can focus our attention first in the decreasing order of the standardized coefficients per group of species. In 5 of the 7 cases, those for groups “All”, “Archaea”, “Eukarya”, animal (“An”) and fungi (“Fu”), the higher standardized coefficient corresponds to *Q*_1_ (black squares), showing that hydrophobic *aa*, encoded with an U base at position *P* 2 of the codons (see Table 2) have a significant larger mean that the *aa* in the other three classes; i.e., in Archaea and, within eukaryotes in animals and plants hydrophobic *aa* have the largest average CUB values, making by association the collections of “All” and “Eukarya” present also *Q*_1_ as the higher coefficients.

The fact that the group of hydrophobic *aa* presents the largest average of CUB values in Archaea, animals and plants (largest *Q*_1_’s in Fig 4) have a rational mechanistic explanation by the importance of the hydrophobic effect, which states that hydrophobic *aa* are usually sequestered away from the aqueous environment into the internal core of a folded protein [17]. In that paper Kyte and Doolittle established that the free energy change associated with transferring an amino acid side chain from water to a non-polar solvent determines its likelihood of being on the surface of the folded protein. Also in [10] Saier mentions that these hydrophobic assignments were likely among the first established in the primordial code to allow for the evolution of the first stable globular proteins.

Interestingly, in Fig 4 Bacteria and plants (“Pl”) are exceptional by the fact that the largest coefficient *Q_i_* is not *Q*_1_, but *Q*_4_ and *Q*_2_, respectively. In bacteria the largest coefficient, *Q*_4_, which contains *aa* with the base G in *P* 2, groups semi-polar *aa* plus Arg, and the corresponding value of the standardized coefficient (shown as a blue diamond in the plot) is significantly larger than the set formed by *aa* in the group *{Q*_1_ *− Q*_2_*}*, which were not significant in the Tukey test. Also intriguing is the fact that in plants the largest coefficient is *Q*_2_, which groups semi-polar *aa* and which codons contains the C base in *P* 2 (red circle in the plot).

Overall –in the “All” group of species, we see that almost half of the average CUB variance is explained by the model of Saier classes (*≈* 47%, shown in the upper part of the “All” column in Fig 4). That percentage is *≈* 44% for fungi, but always larger than 50% for the other groups, reaching its maximum of almost 70% (*≈* 69%) for Archaea. See also **S2.3** in S2 Text.

In summary, from Fig 4 we can conclude that the Saier classification of *aa* is a very important factor influencing the average of CUB values, and, at least in the case of *Q*_1_, we can link the results with a putative molecular explanation (See also **S1.3** in S2 Text for details).

### Effects of the number of codons (***n.cod***) on the average CUB values

The number of codons, *n.cod*, for *aa* with more than one codon in the genetic code (see Table 2), can take values 2, 3, 4 or 6 and, by including the cases for the “Stop” codon, it seems reasonable that this variable could be having an effect on the average CUB values. The rational to think that *n.cod* could have effect on CUB is that a larger value of *n.cod* gives more “degrees of freedom” to the CUB values, even when this coefficient is restricted to values between zero and one. As an example, take the four hydrophobic *aa* (in *Q*_1_, Table 2); of those Leu and Phe are encoded by 6 and 2 different codons, respectively, thus clearly Leu positions within proteins can vary more freely than the ones for Phe and consequently CUB values for Leu could take more extreme values than the ones for Phe during populations bottle necks.

To test the hypothesis that *n.cod* influence average CUB values we can use two alternative parameterizations. First we will consider models including an intercept term, say CUB = ̂*β*_0_ + ̂*β*_1_*n.cod*. Fig 5 shows as examples the three models that explain the largest proportions of CUB variance, selected from the models fitted in the 7 possible groups of species (groups as in the *X*-axis in Fig 4).

**Fig 5.**
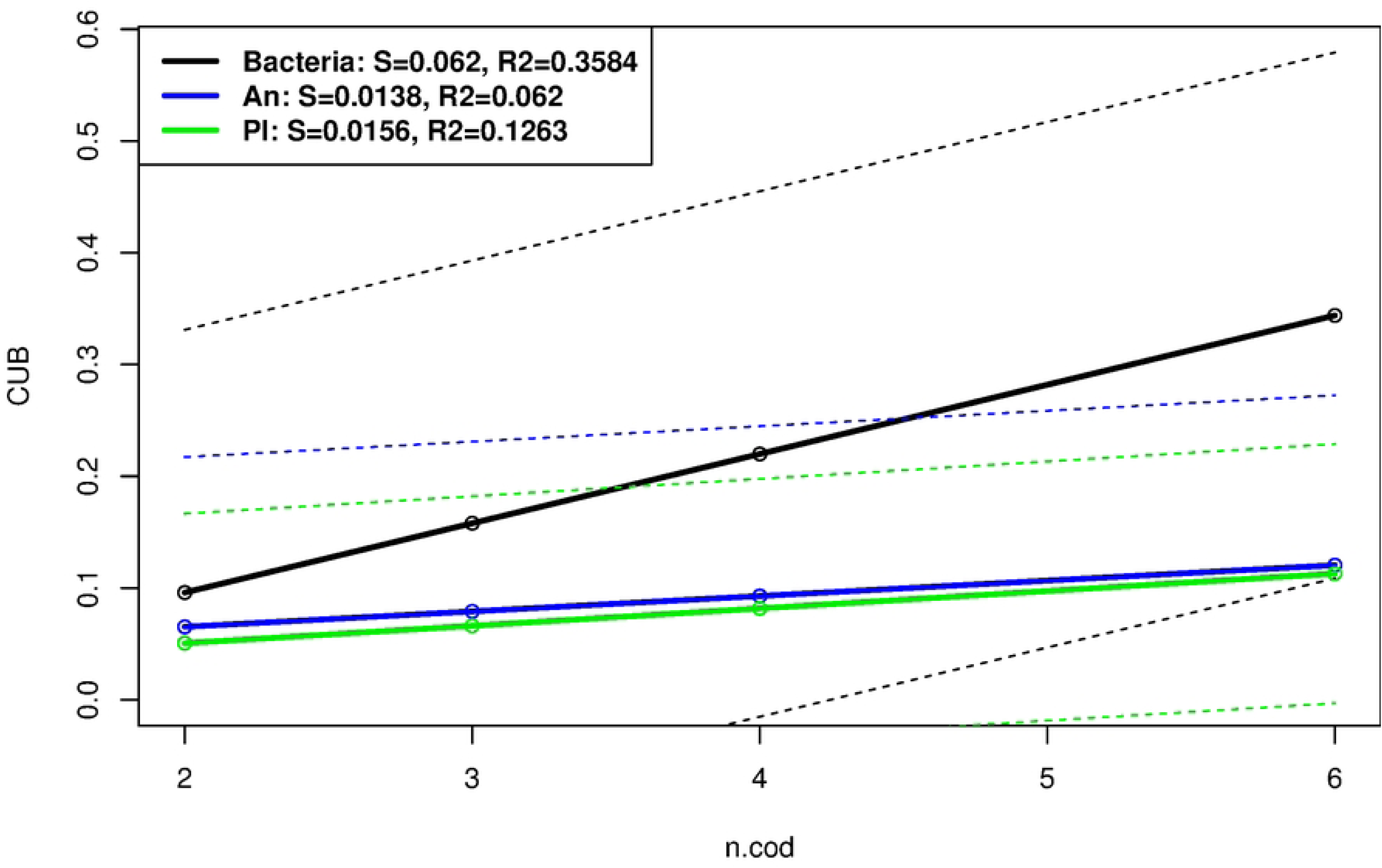
Examples of models. CUB = ̂*β*_0_ + ̂*β*_1_*n.cod*. Continuous lines present the models while dashed lines present the 95% CI for the corresponding models. The slopes and coefficients of determination for the models are presented in the legend as “S” and “R2”, respectively.

In Fig 5 we can see how in bacteria the hypothesis that an increase in the number of codons, *n.cod*, implies an increase in the average CUB value is definitely true; in fact, the *p*-value for the hypothesis that the slope, *β*_1_, in that model is equal to zero is very near zero (*<* 2 *×* 10^−16^; see **S2.4.1** in S2 Text). The estimate of the slope for the bacteria model, *S ≈* 0.062, implies that in average an increase in one unit in *n.cod* “causes” a positive increase of near ten percent, 0.062 *≈* 0.1, in the average CUB value.

In contrast, the increase in average CUB by *n.cod* is much smaller for animals (“An”) and plants (“Pl”), 0.0138 and 0.0156, respectively. However, the 95% Confidence Interval (CI) for the bacteria model –shown as dashed black line in Fig 5, fully includes the models for animals and plants (blue and green continuous lines), indicating that even when the slops of the models are different and in the three cases positive and significant, the models contain a high level of error due to the large CUB variation within the groups. Interestingly the only group for which the slope of the model was not significantly different from zero (*p*-value *≈* 0.09; see **S2.4.1** in S2 Text), correspond to fungi, which was also the group with smaller sample size, only 71 species.

On the other hand, using the alternative parameterization, CUB = *β n.cod*, which excludes the intercept term and thus estimates the means of the effects of 2, 3, 4 and 6 codons on the average CUB values we found that the model including all 1,406 species explains approximately 45% of the CUB variance, but that percentage is only of approximately 36% for fungi and very large, *≈* 73% for bacteria.

The fact that *n.cub* has a larger effect over CUB in bacteria than in all the other groups of species could be influenced by the large population size and small generation time in that group of organisms. That speculation is also backed up by the fact that in Archaea –which also have large population sizes and small generation times, the approximate percentage of CUB variance is also large, *≈* 60%; however, that percentage is slightly larger than 60%, *≈* 60.2% in plants, which in general do not share with bacteria or Archaea the large population sizes and small generation times (see **S2.4.1** in S2 Text for details).

### Effect of ***aa*** volume on average CUB

The variable *vWvol*, the van der Waals radii of *aa*, is measured in cubic Armstrongs (Å^3^), and it represents the total volume occupied by the atoms of each *aa*, a parameter that is crucial for modeling protein folding. The rational to look for effects of the *aa* volume on the average CUB value is that larger *aa* could have a larger average CUB than smaller ones due to structural constrains within proteins.

In the 18 *aa* coded by more than one codon *vWvol* varies from a minimum of 48 for Gly up to a maximum of 148 Å^3^ for Arg, i.e., there is a difference of approximately 3 times between the largest and smallest *aa*. Among the 18 *aa*, *vWvol* takes only 16 different values, due to the fact that Ile and Leu have both a value of 124 Å^3^, while Phe and Lys share a volume of 135 Å^3^.

Fig 6 presents the 3 models CUB = ̂*β*_0_ + ̂*β*_1_*vW vol* for the groups of species in which the models explained more than 1.2% of the CUB variance.

**Fig 6.**
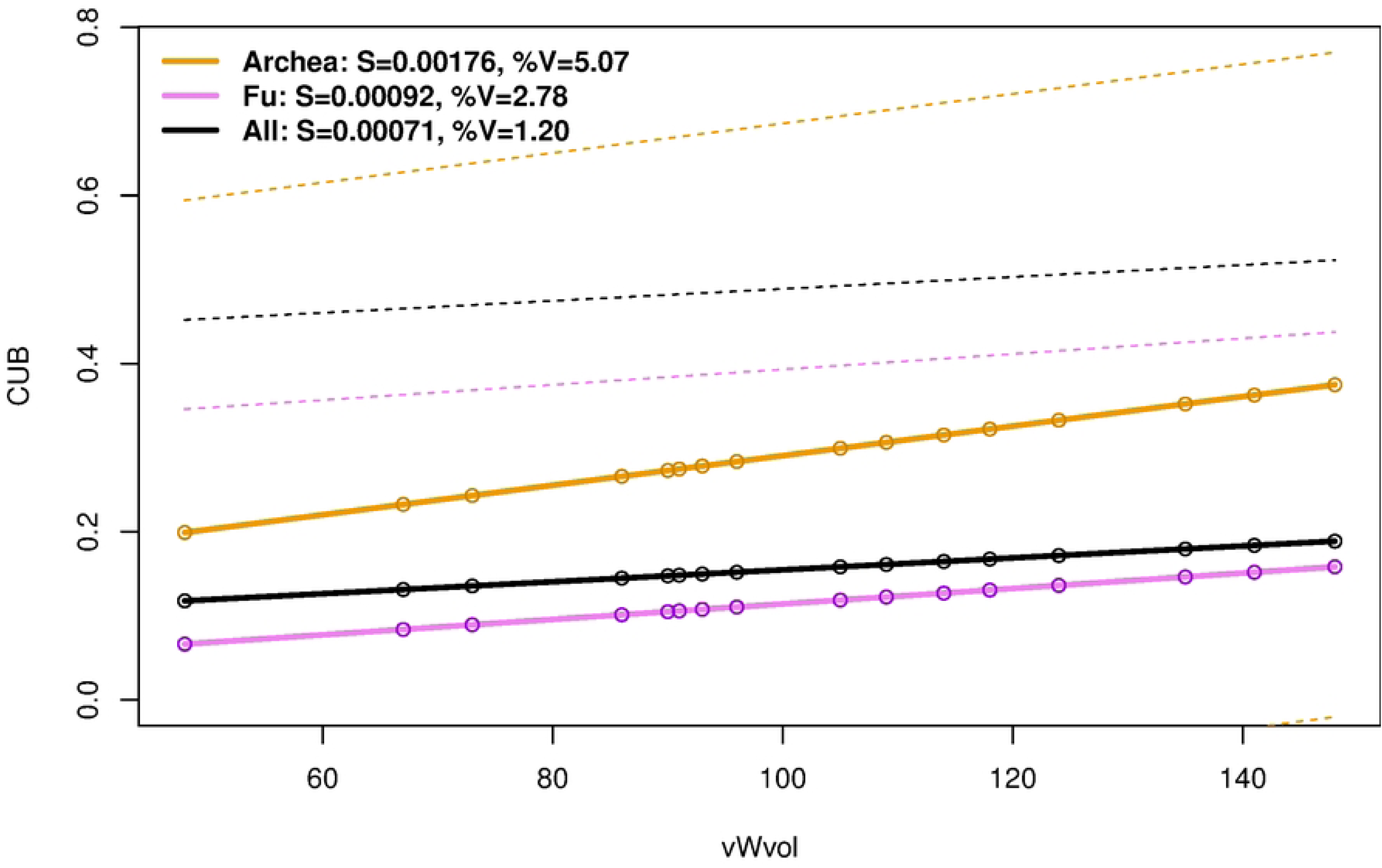
Examples of models. CUB = ̂*β*_0_ + ̂*β*_1_*vW vol*. Continuous lines present the models while dashed lines present the 95% upper CI for the corresponding models. The slopes, “S”, and percentages of explained variance and, “%V”, are in legend.

In Fig 6 we can see that the three models shown have a positive but small slope (̂*β*_1_) that for the model of the whole set of species (model All in the figure) takes a value 0.00071 while for the case of Archean is approximately 2.5 times larger, taking a value of 0.00176. Even when in all seven groups of species the corresponding models have large statistical significance, the small proportion of CUB variance explained make them biologically irrelevant; thus we can conclude that the volume of the *aa* does not have a substantial effect on our variable of interest (See **S2.4.2** in S2 Text for details).

### Effect of ***aa*** hydrophobicity on CUB by group of species

We have seen before, when analyzing the effect of Saier classes on average CUB values, how hydrophobicity is a relevant factor which affects CUB. In here we study hydrophobicity independently from other factors by taking into account only the sets of the 11 *aa* which are coded by at least 2 codons and also are classified with an hydrophobic or hydrophilic class, say the sets i=*{*Asn, Asp, Gln, Glu, His, Lys, Tyr*}*, of the the 7 hydrophilic *aa* and o=*{*Ile, Leu, Phe, Val*}*, the set of the 4 hydrophobic ones.

By fitting the models CUB = *I_h_β*, where *I_h_* is a dummy variable indicating if the *aa* is hydrophobic or hydrophilic for each group of species we obtained highly significant models in which the *p*-value for the hypothesis *β* = 0 was negligible (*p*-value *≈* 0 in all cases). Those models explained a minimum of 44% of the CUB variance in the group of fungi (Fu), and a maximum of 69% of the CUB variance in Archaea. The overall percentage of explained variance in the group including all species was of 45%, demonstrating that *aa* hydrophobicity is highly relevant factor affecting the average CUB values in all species (See **S2.4.3** in S2 Text for details).

Fig 7 presents the distributions, as box plots, of the CUB value by hydrophobicity and species group.

**Fig 7.**
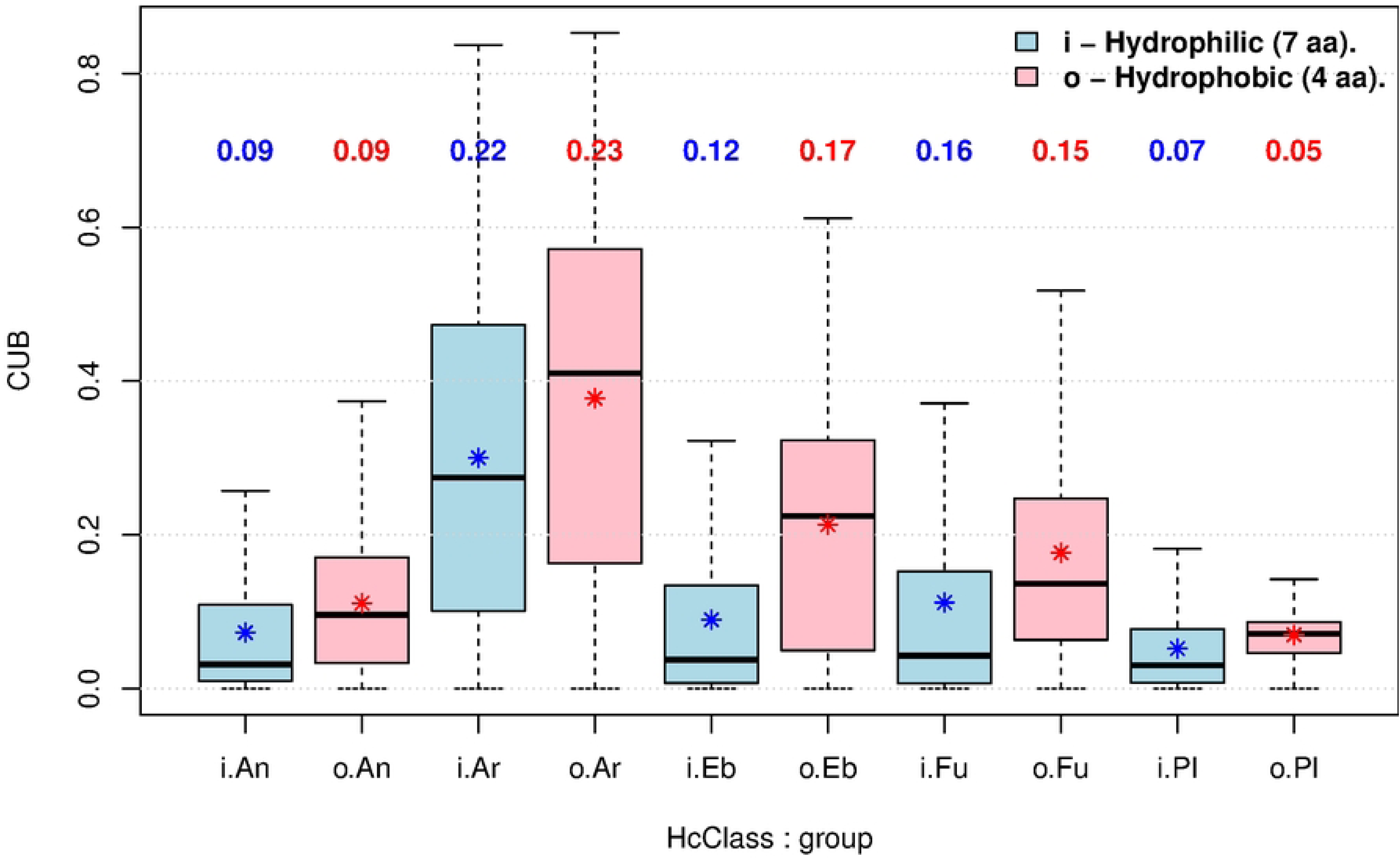
Distributions of CUB by hydrophobicity and species group. The colored numbers at *Y* -axis = 0.7 give the standard deviations (*S*^^^) of the corresponding distributions. Asterisks signal the means (averages) of CUB in each box plot.

In Fig 7 the first notorious fact is that in all main groups of species the mean of CUB values is larger for hydrophobic (red asterisks) than for hydrophilic *aa* (blue asterisks). This is in line with the importance of the hydrophobic effect [17], that as commented above, implies that hydrophobic *aa* tend to be into the internal core of the folded proteins, and thus changes from hydrophobic to hydrophilic *aa* will be generally very deleterious, which in turn is likely to produce the observed difference. In other words, hydrophobic *aa* tend to keep a larger CUB value during evolution than hydrophilic ones because selection forces are likely to be stronger in the former group than in the latter.

The differences of the mean CUB values between the sets of hydrophobic and hydrophilic *aa* go from a minimum of 0.01760 in plants (Pl) up to a maximum of 0.12378 in Bacteria, with an overall change of 0.0575 in all the species (group not shown in Fig 7). In fact, those differences are larger on microbes (Archaea with a value of 0.0775 and Bacteria with a value of 0.1238), than in eucaryotes (An, Pl and Fu), which have a value of 0.0373 for that difference. That could be due to the fact that in general microbes have larger population sizes and smaller generation times than eucaryotes, making selection factors more immediate and efficient in the former group, but that hypothesis does not fit for plants, which have the smallest difference.

### Effect of the aliphatic / aromatic class on CUB by group of species

In Table 2 we can see that there are 7 *aa* classified as aliphatic, the set l= *{*Ala, Ile, Leu, Val*}*, while the set of aromatic ones is r= *{*His, Phe, Tyr*}*.

The distinction between aliphatic and aromatic *aa* is central to protein stability, folding, and molecular recognition. While both groups are predominantly hydrophobic and tend to be buried within the protein core, their specific roles differ due to the unique electronic properties of aromatic rings [18].

To detect the unique effect of the aliphatic / aromatic classification on the average CUB value we fitted models CUB = *β*_0_ + *β*_1_*A*, where the value of *A* is equal to -1/4 if the *aa* is aliphatic while it takes the value of 1/3 for the cases where the *aa* is aromatic, being undetermined (NA) for *aa* out of the relevant sets. With this parameterization the model captures the influence of the aliphatic / aromatic class on CUB, given that if the CUB value is the same for aliphatic and aromatic *aa* the value of ̂*β*_1_ will be zero.

For all groups of species, except for Archaea, the hypothesis: *β*_1_ = 0 for models CUB = *β*_0_ + *β*_1_*A* was rejected with negligible *p*-values very near to zero. For Archaea the *p*-value was 0.257, indicating that for that group the aliphatic / aromatic classification was not affecting the average CUB (See **S2.4.4** in S2 Text for details).

Fig 8 presents the distributions, as box plots, of CUB values segregated by aliphatic / aromatic class and main group of species.

**Fig 8.**
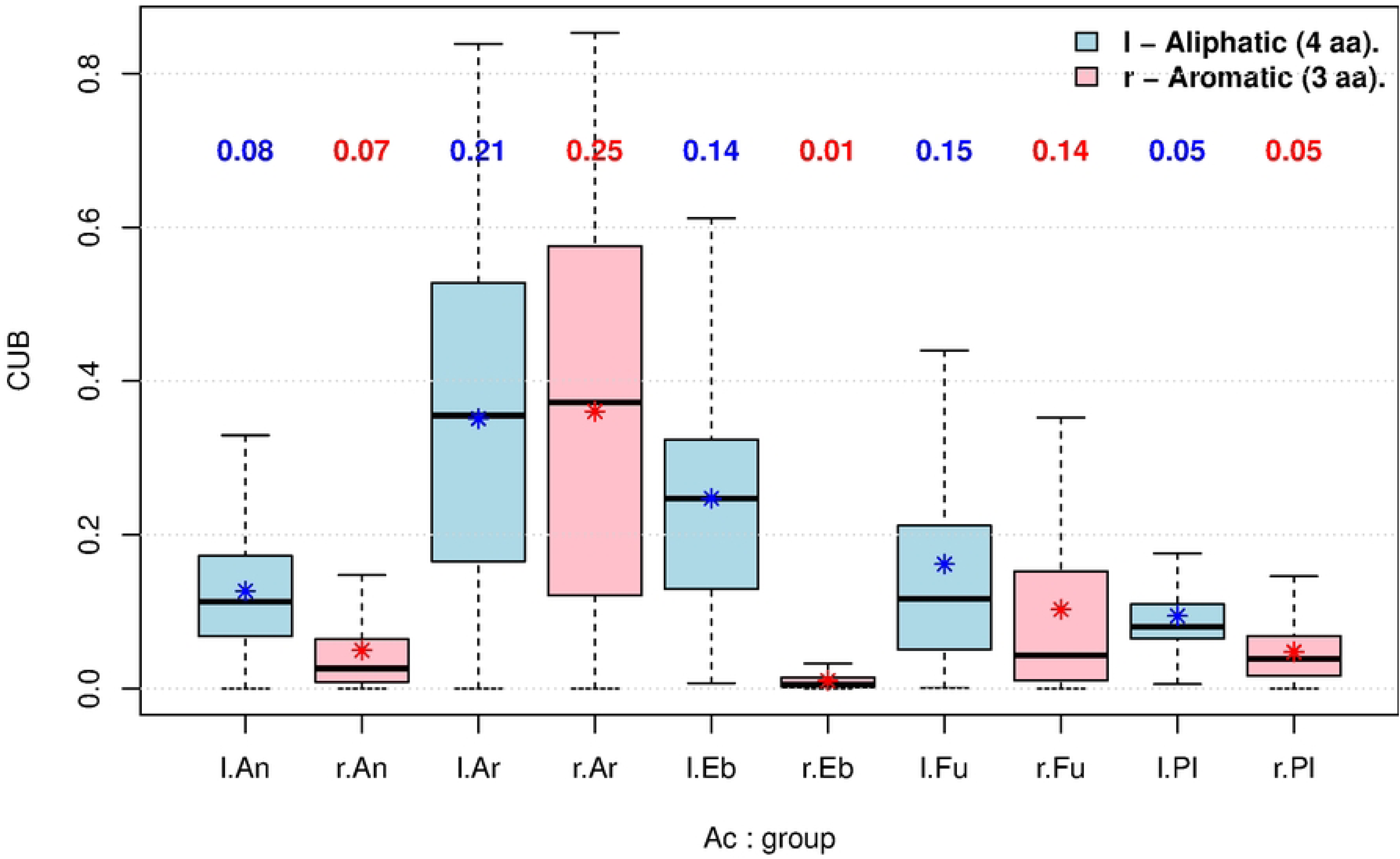
Distributions of CUB for “Aliphatic / Aromatic” class and group of species. The color numbers at *Y* -axis = 0.7 give the standard deviations (*S*^^^) of the corresponding distributions. Asterisks signal the means (averages) of CUB in each box plot.

In Fig 8 we can see that in all groups, except in Archaea, there are large CUB differences between the aliphatic and aromatic groups. In all these cases the mean CUB value is larger for aliphatic than for aromatic *aa*, and the largest difference occurs in bacteria (group “Eb”), where the distribution of CUB in aromatic *aa* (bar with label “r.Eb” in Fig 8) is very close to zero and with a negligible dispersion (standard deviation *≈* 01).

Employing a different parameterization, say, fitting the models CUB = *β*_1_*A_l_* + *β*_2_*A_r_*, where *A_l_* and *A_r_* are dummy variables which are equal to 1 if the *aa* are aliphatic and aromatic, respectively, and 0 otherwise, we effectively segregate the effect of each one of the two classes on CUB, without taking into account the possible interaction between the classes. Given that those models do not include an intercept term, taking into account only the absolute effects of aliphatic –by *β*_1_ and aromatic –by *β*_2_ *aa*, the proportion of CUB variance explained by those models is large and statistically significant for all groups of species, being 48% for the set of all species and very large for microbes, 71 and 75% for Archaea and Bacteria, respectively. While models CUB = *β*_0_ + *β*_1_*A* take into account only the weighted difference in average CUB between the classes, which as seen in Fig 8 is not significant for Archaea, models CUB = *β*_1_*A_l_* + *β*_2_*A_r_*separate the effects of each one of the two classes. Estimates of these models for each group of species are presented within **S2.4.4** in S2 Text includes a more detailed discussion of the effects of the aliphatic / aromatic effects on CUB.

### Effects of dissociation constants ***pK*_1_** and ***pK*_2_** on the average CUB values

Table 2 shows the numeric values of the dissociation constants *pK*_1_ and *pK*_2_ for the 18 *aa* encoded by at least two codons. The dissociation constants *pK*_1_ (for the *α*-carboxyl group) and *pK*_2_ (for the *α*-amino group) are fundamental to the chemical identity of amino acids, determining their ionization state and net charge at a fixed pH [19]. Given this fact, we fitted models of average CUB values per group of species to raw and normalized versions of the *pK*’s. From those models we observed that *pK*_1_ is significantly (*p*-value *<* 0.01) with CUB, while *pK*_2_ results significant for CUB values in eukaryotes, but not in Archaea (*p*-value = 0.2458), and only slightly significant (*p*-value = 0.0493) in Bacteria (See **S2.4.5** in S2 Text for details).

### Percentages of explained CUB variance in individual models

In the previous section we obtained the relations of average CUB values with different *aa* characteristics per groups of species. However, given that we have a variable number of species per group (see Table 1), what is true for the average CUB in a given group of species could be inaccurate for some (or many) of the species in a given group. To solve this problem and obtain a clearer panorama of the effect of the *aa* characteristics on individual species, we fitted individual models for CUB and *aa* characteristic in each one of the 1,406 species studied and which belong to only one of the main groups, say, animals (An), bacteria (Eb), Archaea (Ar), fungi (Fu) or plant (Pl); see Table 1.

The parameterization of those models per specie excluded the intercept term and included variables for each one of the relevant classes within each *aa* characteristic. For example, for the variable number of codons, *n.cod*, the model includes a single coefficient, say *β*, and the model for a given species is CUB = *β n.cod*, where *n.cod* takes values 2, 3, 4 or 6, i.e., four different values which are assigned to each one of the 19 *aa* –including the “Stop” signal, thus the model for a given species includes 19 CUB values. In contrast, for Saier classes, the model for a single species is CUB = *β_i_ Q_i_*, where each *β_i_*measures the effect of the class *Q_i_*, and the model for a given species is valid for only 18 *aa* –because the “Stop” codons are not *aa* and thus the characteristics associated with the four Saier classes does not make sense, and thus the model fits 18 CUB values in a given species.

Because it will be very difficult to analyze in detail the 1,406 models for each one of the species in each one of the *aa* characteristics, we use the distributions of the percentage of CUB variance explained by the models as a proxy to judge the relative importance of each *aa* characteristic on each group of species (See **S2.5** in S2 Text for details). Fig 9 presents those distributions.

**Fig 9.**
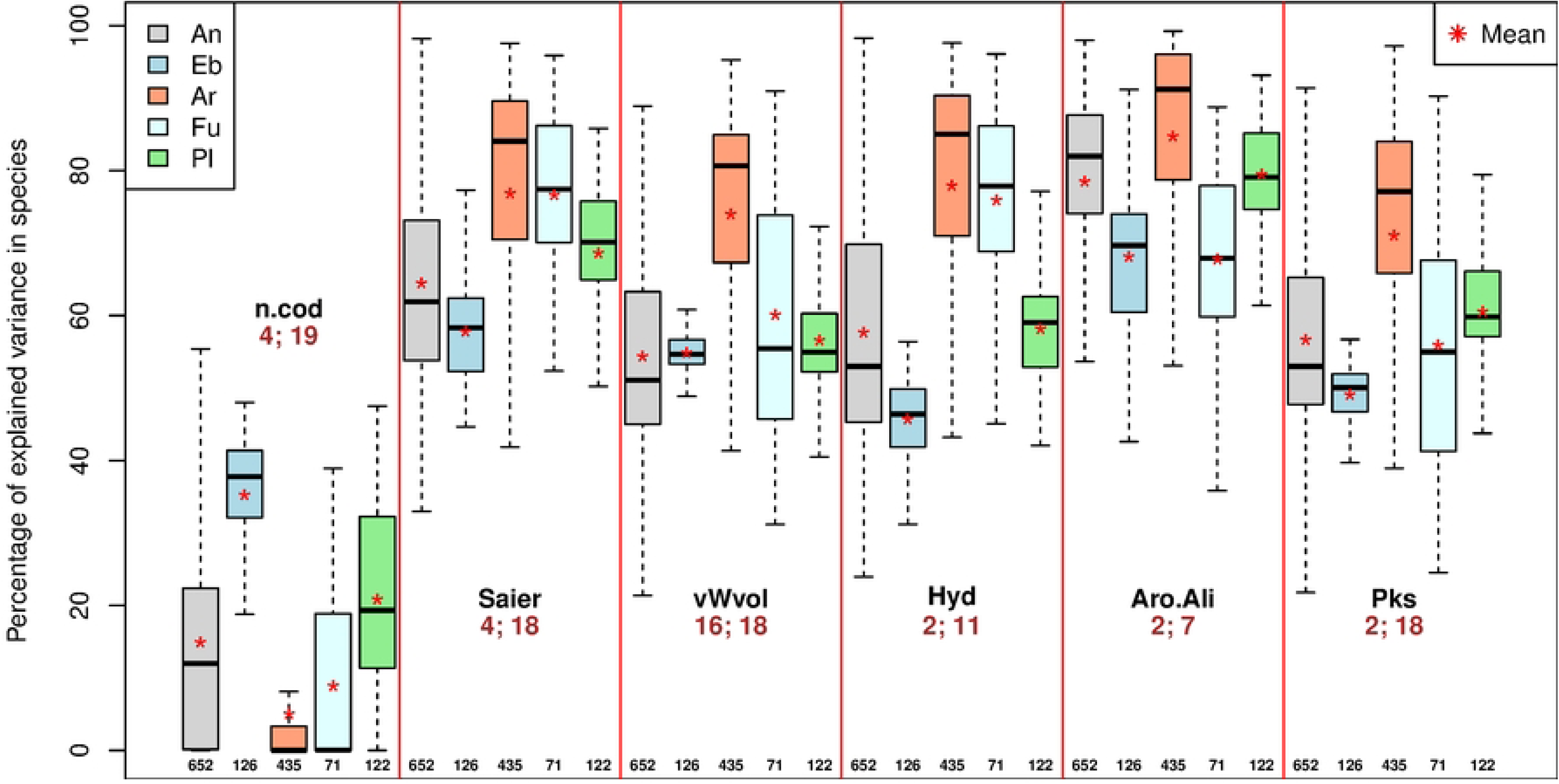
Percentages of CUB variance by factor in models by species. Distributions as box plots of percentages of explained variances, *V* % = 100 *×* 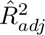, , of models fitted for individual species for each one of the 6 variables studied (*aa* characteristics: *n.cod* - number of codons, Saier - *P* 2 class, *vWvol* - *aa* volume, Hyd - *aa* hydrophobicity, Aro.Ali - Aromatic/Aliphatic class and pKs - Dissociation constants). Below the name of each variable the brown numbers give the number of classes and *aa* in each one of the models, respectively. The number of species (models) is presented for each variable at the bottom of the corresponding distribution and no points out of the whiskers (outliers) are shown.

Even when in Fig 9 we do not have the *p*-values associated with the models, we can judge the biological relevance of the individual models by the percentages of CUB variance explained, which is given in the *Y* -axis; for example, models which explain less than 10% of the CUB variance will be likely irrelevant, while models explaining at least 60% of that variance are much more interesting.

The first distinguishing feature in Fig 9 is that *n.cod* models explains less proportion of the CUB variance in models for individual species. In fact, only in bacteria (Eb group) the median of the distribution is close to 40% –on line with the results presented in Fig 5, where we saw that the slope for average CUB as function of *n.cod* in bacteria is large. Nevertheless, we will see in Table 3 that there are outlier species which present a higher %V in other groups for *n.cod*. Also interesting in the distributions of %V for models of CUB by *n.cod* is the fact that the distribution for Archaea (Ar) is extremely compact and close to the lower bound, %V = 0. Even when arguing that CUB and *n.cod* could have a causal relation is dubious, what is clear is that the distributions of %V from individual models are highly different between bacteria and archaea.

**Table 3.** Species with maximum CUB explained variance (%*V* ) by variable and group in individual models.

| **Variable: $n_{cod}$ (number of codons per $aa$)** |  |  |  |  |
| --- | --- | --- | --- | --- |
| Group | $n_{mod}$ | %V | Key | Species (comment) |
| An | 652 | 58.11 | An96 | *Bufo bufo* (European toad) |
| Eb | 126 | 48.00 | Eb69 | *Klebsiella aerogenes* (Pathogenic) |
| Ar | 435 | 50.27 | Ar231 | *Methanobrevibacter thaueri* (Methanogenic Archaea) |
| Fu | 71 | 38.89 | Fu50 | *Serpula lacrymans* (Causes dry rot) |
| Pl | 122 | 47.49 | Pl65 | *Morus notabilis* (Chinese mulberry) |
| **Variable: Saier class (4 quadrants)** |  |  |  |  |
| Group | $n_{mod}$ | %V | Key | Species (comment) |
| An | 652 | 98.20 | An512 | *Pollicipes pollicipes* (A goose barnacle; Arthropoda) |
| Eb | 126 | 77.31 | Eb67 | *Franconibacter helveticus* (A bacteria of the Cronobacter genus) |
| Ar | 435 | 97.57 | Ar233 | *Methanobrevibacter wolinii* (A methanogen archaeon) |
| Fu | 71 | 95.87 | Fu44 | *Rhodotorula graminis* (Fungi in the class Microbotryomycetes) |
| Pl | 122 | 95.41 | Pl122 | *Chlamydomonas reinhardtii* (Single-cell green alga) |
| **Variable: $vWvo$ ($aa$ volume; van der Waals radii)** |  |  |  |  |
| Group | $n_{mod}$ | %V | Key | Species (comment) |
| An | 652 | 93.39 | An512 | *Pollicipes pollicipes* (A goose barnacle; Arthropoda) |
| Eb | 126 | 64.87 | Eb113 | *Salmonella bongori* (Pathogenic bacterium) |
| Ar | 435 | 95.27 | Ar174 | *Halorussus rarus* (Halophilic archaeal) |
| Fu | 71 | 90.97 | Fu12 | *Cutaneotrichosporon oleaginosum* (Family Trichosporonaceae) |
| Pl | 122 | 90.39 | Pl122 | *Chlamydomonas reinhardtii* (Single-cell green alga) |
| **Variable: Hydrophobicity** |  |  |  |  |
| Group | $n_{mod}$ | %V | Key | Species (comment) |
| An | 652 | 98.26 | An512 | *Pollicipes pollicipes* (A goose barnacle; Arthropoda) |
| Eb | 126 | 64.26 | Eb105 | *Plesiomonas shigelloides* (Associated with diarrhea) |
| Ar | 435 | 97.60 | Ar311 | *Methanothermus fervidus* (Methanogen and thermophilic) |
| Fu | 71 | 96.09 | Fu44 | *Rhodotorula graminis* (Fungi in the class Microbotryomycetes) |
| Pl | 122 | 94.87 | Pl122 | *Chlamydomonas reinhardtii* (Single-cell green alga) |
| **Variable: Aromatic / Aliphatic** |  |  |  |  |
| Group | $n_{mod}$ | %V | Key | Species (comment) |
| An | 652 | 97.98 | An587 | *Strongyloides ratti* (Parasitic nematode) |
| Eb | 126 | 91.15 | Eb37 | *Cronobacter universalis* (Facultatively anaerobic) |
| Ar | 435 | 99.24 | Ar120 | *Halopelagius longus* (Thermotolerant, extremely halophilic archaeon) |
| Fu | 71 | 88.78 | Fu50 | *Serpula lacrymans* (Causes dry rot) |
| Pl | 122 | 93.15 | Pl65 | *Morus notabilis* (Chinese mulberry) |
| **Variable: Dissociation constants ($pK_1$ and $pK_2$)** |  |  |  |  |
| Group | $n_{mod}$ | %V | Key | Species (comment) |
| An | 652 | 97.14 | An587 | *Strongyloides art* (Parasitic nematode) |
| Eb | 126 | 56.71 | Eb67 | *Franconibacter helveticus* (A bacteria of the Cronobacter genus) |
| Ar | 435 | 97.18 | Ar21 | *Methanobrevibacter arboriphilus* (Strictly anaerobic archaea) |
| Fu | 71 | 90.27 | Fu44 | *Rhodotorula graminis* (Fungi in the class Microbotryomycetes) |
| Pl | 122 | 84.98 | Pl122 | *Chlamydomonas reinhardtii* (Single-cell green alga) |

$n_{mod}$ : Number of models from which the one with maximum explained variance (%V) was selected (equal to the number of species in the corresponding group). %V: Maximum in percentage of the determination coefficient, $\hat{R}_{adj}^2$ , among the $n_{mod}$ fitted. Key: Unique identifier of the species in the data set.

The second interesting characteristic in Fig 9 is that the distributions for the variables Saier, *vWvol*, Hyd, Aro.Ali and *Pk*s are relatively alike in that the interquartile range (the box in the box plots) in all those distributions is higher for Archaea than for the other 4 species groups, and the same happens with the corresponding means and medians. This implies that, with the intriguing exception of *n.cod*, the CUB values are more affected by the *aa* characteristics studied in Archaea than in the other species’ groups. On the other hand, the likeness of the distributions for Saier, *vWvol*, Hyd, Aro.Ali and *Pk*s within the different groups of species are in part due to the interdependence that exist between the variables.

The more relevant of the variables studied in relation with CUB is the classification by Saier classes, which involves both, genetic—the base at the second position of the codon, as well as biochemical characteristics of the *aa*—polarity and hydrophobicity. In Fig 9 for the Saier classes we can see that the decreasing order of means of percentages of explained variances (*Y* -axis) are approximately 77, 77, 69, 64 and 58, corresponding to Ar Fu Pl An and Eb, respectively, while the medians are approximately 84, 77, 70, 62 and 58 with the same order of groups, Ar Fu Pl An and Eb, respectively. This implies that, on average, models for individual species explain a largest percentage of explained variance for Archaea (Ar *≈* 77%) than for bacteria (Eb *≈* 58%), while the mean percentages for eukaryotes are intermediate (Fu also *≈* 77%, but with much smaller median than Ar, Pl *≈* 69 and An *≈* 64%).

Judging the dispersion of the distributions of the percentages of explained variances of the CUB models in Saier classes (shown in Fig 9) by the standard deviations of those values we find that the order from more to less variable distributions correspond to the groups Ar (*S*^^^ *≈* 18%), An (*S*^^^ *≈* %12), Fu (*S*^^^ *≈* 11%), Pl (*S*^^^ *≈* 9%) and Eb (*S*^^^ *≈* 7%); thus, the percentage of explained variance presents, in general, large variation within groups of species, implying that there is not a fully reliable rule to predict how much CUB variance will be explained by Saier classes in a particular organism.

Nevertheless, we can examine the minimum of explained variance of the CUB models in Saier classes per group of species. Those minima are 52% for Fu, 45% for Eb, 41% for Pl, 33% for An and 22% for Ar. Those minima are not included within the whiskers of the Saier distributions in Fig 9, because we omit outliers, however we can be reasonably sure that, even in the worst of the cases, Saier classes explain a substantial proportion of the CUB variance.

On the other hand, we can see in which species each one of the variables reached the maximum of CUB explained variance. Table 3 presents the species with maximum CUB explained variance (%*V* ) by variable and group in individual models.

Even when presenting in Table 3 only the species which reached the maximum of explained CUB variance for each one of the variables studied distorts the perception of “how good” the individual models could be, also exemplifies the extreme cases where a very large part of the CUB variance is explained by an *aa* characteristics. In general, the points corresponding to the species in Table 3 are not shown in Fig 9, because they are outliers for the corresponding distributions.

In the section of Table 3 corresponding to the number of codons (*n.cod*) per *aa*, we see that the smallest %V, 38.89, corresponds to a fungus, *Serpula lacrymans* which causes dry rot in wood, while the largest %V, 58.11, happens in the European toad *Bufo bufo*. Thus, even when in Fig 9 we saw that *n.cod* is the variable which explains a smaller fraction of the CUB variances, there are exceptions in species where a substantial fraction of those variances correlates with the number of codons that determine the *aa*.

Much higher values of %V are shown in Table 3 for the variable Saier class, related with both, the second base in the codon as well as *aa* polarity and hydrophobicity. In that case the smallest %V, is very high, 77.31, and corresponds to a facultative anaerobe bacteria of the Cronobacter genus, *Franconibacter helveticus*, while the largest %V, 98.20, occurs for *Pollicipes pollicipes*, an edible goose barnacle, of the phylum Arthropoda. It is interesting that for Archea and eucaryotes we find species in which more than 95% of the CUB variance is explained by the Saier class; for example in plants we have the single-cell green alga *Chlamydomonas reinhardtii* in which approximately 95.41% of CUB variance is explained by that variable.

For the other 4 variables studied, *aa* volume (*vWvo*), hydrophobicity, Aromatic / Aliphatic classes and dissociation constants (*pK*_1_ and *pK*_2_), Table 3 shows that the %V explained is always *>* 56%, and frequently *>* 90%, demonstrating that along the three life domains there are individual species in which the *aa* characteristics are highly correlated with CUB.

Even then Table 3 has 5 *×* 6 = 30 rows, corresponding to the combinations of main groups per variables, it includes only 19 different species, given that the same organism was found repeatedly as having the maximum %V in more than one variable. For example, the single-cell green alga *Chlamydomonas reinhardtii* appears 4 times in Table 3, because it had the maximum %V for variables Saier, *vWvol*, hydrophobicity and *P_k_*s. That species has an average of %V of 91.41% over the 4 variables, being a clear outlier for CUB explained variance among all the species studied. Interestingly, *C. reinhardtii* was also found to be an outlier, nor only for CUB, but also for all informational measures related with the genetic code [9].

Other species that were found more repeated 3 times in Table 3 are *Pollicipes pollicipes* –for variables Saier, *vWvol* and Hydrophobicity and with an average %V = 96.62 and *Rhodotorula graminis* –for variables Saier, Hydrophobicity and *pK*’s and with an average %V = 94.08. Also, other 4 species were found twice in Table 3 (See **S2.5** in S2 Text for details).

Using linear models, we have demonstrated relations between CUB and physicochemical *aa* characteristics in a sample that includes organisms from the three domains of life. However, the mere observation of these relations is insufficient to disentangle the history of genomic changes during the evolution of a lineage; we are looking through a very narrow window which shows only how different clades have developed particular “accents” in the use of the genetic code.

The current integrated theory of molecular evolution [20] underlines the relevance of the drift-barrier hypothesis [21, 22]. This hypothesis posits that, like all traits, the mutation rate is subject to evolutionary modification; thus, natural selection can only reduce mutation rates or optimize codon usage to the point where the selective advantage outweighs the power of random genetic drift. Evolutionary rates are not merely products of selection, but are proximately governed by metabolic and genomic machinery (such as DNA repair and generation time), which are themselves under selective pressure. This explains why CUB is often more pronounced in species with large effective population sizes, as selection for translational efficiency can overcome the “noise” produced by genetic drift.

### Dendrograms from CUB values

In the margins of Fig 2 we observed and discussed the dendrograms obtained by the medians of CUB values for the main groups of species in the *X*-axis and for the *aa* in the *Y* -axis. In particular, the dendrogram for the main groups of species in that figure is a phenogram [23], i.e., a digram clustering group of species by similarities in one aspect of the genomic dialects rather than a reconstructed phylogeny.

Section **S2.6** in S2 Text presents and briefly discuss the similarities in CUB values for the *aa* in the full set of the 1,434 species studied.

On the other hand, we wanted to investigate the topologies of dendrograms based on CUB values and obtained from random samples of 5 species, one from each one of the 5 main groups (Ar, Eb, An, Pl and Fu; see Table 1), to estimate the relative frequency in which the true topology between the groups, say, *T_t_* = *{*Eb, *{*Ar, *{*Pl, *{*Fu, An*}}}}* [15], was recovered from dendrograms constructed from those random samples, as well as to estimate the frequency in which the possible clusters of species appeared in those dendrograms.

With this aim, we obtained *b* = 100, 000 random samples of 5 species, one from each one of the 5 main groups, and constructed the *b* dendrograms from the CUB values of the selected species, evaluating the frequency in which each one of the different *n_t_* = 105 topologies appeared. The topology of that appeared more frequently among the *b* dendrograms was

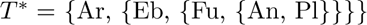

which show up in 18,112 of the *b* cases, i.e., in a relative frequency of approximately 18.11% of the random dendrogram. In contrast, only in 1,767 of the *b* cases, i.e., in approximately 1.77% of the cases the topology of the observed dendrograms was equal to the true topology *T_t_*, demonstrating that dendrograms from random samples of the species studied rarely coincided with the true topology, *T_t_*. This implies that CUB values are not a good proxy for phylogenetic inference.

Section **S2.7** in S2 Text presents a detailed analysis of the topologies of the *b* random dendrograms obtained and discuss the implications of the differences between the true phylogenetic relations between the groups of species and the ones observed in dendrograms from random samples, while section **S2.8** in S2 Text presents and discusses the set of distances in the CUB space.

## Conclusion

The systematic analysis of CUB through the lens of amino acid physicochemical properties confirms that the genetic code is a dynamic informational system. Our results demonstrate that the choice of synonymous codons is not a neutral process but is strictly partitioned by the biochemical identity of the residues they encode. The strong influence of the second codon position (*P* 2) and hydrophobicity on CUB values across all domains suggests that these “genomic dialects” are rooted in primordial error–minimization strategies and the thermodynamic requirements of protein folding.

Furthermore, the high variance in CUB explained by individual species models underscores the diversity of informational accents. While microbes generally exhibit more pronounced biases due to large effective population sizes and selection for translational efficiency, the consistency of these patterns across eukaryotes points to a universal scaling of molecular evolution governed by the drift–barrier hypothesis. By distinguishing between phenetic similarity (dendrograms) and phylogenetic history, we provide a clearer perspective on how clades have developed unique “accents” while maintaining the near–optimal informational potential of the universal genetic code.

## Supporting information

**S1 Dataset. CUB values.** Columns are “Species” and “Key” followed by the *aa* abbreviations. For each combination of row and *aa* the data are CUB values. (ZIP).

**S2 Text. Supporting text.** Given the extensive nature of our results, all supporting text, figures, and tables are compiled in the “S2 Text.pdf” file. These supporting information are referenced in the main text using bold faced labels. (PDF).

**S3 R. R objects.** All objects containing intermediate and final results were compiled and deposited in Zenodo at: https://zenodo.org/records/19609962.

## References

1. Ji S. Isomorphism between cell and human languages: molecular biological, bioinformatic and linguistic implications. BioSystems. 1997;44(1):17–39. doi:10.1016/S0303-2647(97)00039-7.

2. Jakobson R. Linguistics in Relation to Other Sciences. In: Selected Writings, Vol. 2: Word and Language. The Hague: Mouton; 1971. p. 655–96. Discusses the formal similarities between the genetic code and the structural laws of human language.

3. Barbieri M. The Organic Codes: An Introduction to Semantic Biology. Cambridge: Cambridge University Press; 2003. Introduces the idea of ”organic semantics,” providing a strong foundation for the ”accent” metaphor in CUB.

4. Grantham R, Gautier C, Gouy M, Mercier R, Pave A. Codon catalog usage is a genome strategy modulated for gene expressivity. Nucleic Acids Research. 1980;8(1):r49–62.

5. Ikemura T. Codon usage and tRNA content in unicellular and multicellular organisms. Molecular Biology and Evolution. 1985;2(1):13–34. doi:10.1093/oxfordjournals.molbev.a040335.

6. Sharp PM, Li WH. The codon adaptation index-a measure of directional synonymous codon usage bias, and its potential applications. Nucleic Acids Research. 1987;15(3):1281–95.

7. Wright F. The ‘effective number of codons’ used in a gene. Gene. 1990;87(1):23–9.

8. Martínez O. Shannon.codon: An R package for the analysis of codon frequencies. Zenodo [Internet]. 2025 Jun [cited 2026 March 30];Available from: 10.5281/zenodo.15650136.

9. Martínez O, Reyes-Valdés MH, Ochoa-Alejo N. Sampling informational properties of codon usage through the tree of life. PLOS ONE. 2025;20(11):e0335824. doi:10.1371/journal.pone.0335824.

10. Saier MH. Understanding the Genetic Code. Journal of Bacteriology. 2019;201(15):10.1128/jb.00091-19. Available from: https://journals.asm.org/doi/abs/10.1128/jb.00091-19. arXiv:https://journals.asm.org/doi/pdf/10.1128/jb.00091-19. doi:10.1128/jb.00091-19.

11. Subramanian K, Payne B, Feyertag F, Alvarez-Ponce D. The codon statistics database: a database of codon usage bias. Molecular Biology and Evolution. 2022;39(8):msac157.

12. Wikipedia contributors. Proteinogenic amino acid — Wikipedia, The Free Encyclopedia [Internet]; 2026 [cited 2026 March 30]. Available from: https://en.wikipedia.org/w/index.php?title=Proteinogenic_amino_acid.

13. Martínez O. DendroLikeness: an R package to compare dendrograms [Internet]. Zenodo; 2024 [cited 2026 March 30]. Available from: 10.5281/zenodo.13737570.

14. R Core Team. R: A Language and Environment for Statistical Computing [Internet]; 2025 [cited 2026 March 30]. Available from: https://www.R-project.org/.

15. Williams TA, Foster PG, Cox CJ, Embley TM. An archaeal origin of eukaryotes supports only two primary domains of life. Nature. 2013;504(7479):231–6. doi:10.1038/nature12779.

16. Baldauf SL, Palmer JD. Animals and fungi are each other’s closest relatives: congruent evidence from multiple proteins. Proceedings of the National Academy of Sciences. 1993;90(24):11558–62.

17. Kyte J, Doolittle RF. A simple method for displaying the hydropathic character of a protein. Journal of molecular biology. 1982;157(1):105–32. doi:10.1016/0022-2836(82)90515-0.

18. Makwana KM, Mahalakshmi R. Implications of aromatic–aromatic interactions: From protein structures to peptide models. Protein Science. 2015;24:1920–33. doi:10.1002/pro.2814.

19. Grimsley GR, Scholtz JM, Pace CN. A summary of the measured pK values of the ionizable groups in folded proteins. Protein Science. 2008;18:247–51. doi:10.1002/pro.19.

20. Caron FS, Domingos FMCB. Revisiting the determinants of molecular evolutionary rate variation. Evolutionary Journal of the Linnean Society. 2026 02;5(1):kzag003. Available from: 10.1093/evolinnean/kzag003. arXiv:https://academic.oup.com/evolinnean/article-pdf/5/1/kzag003/66820900/kzag003.pdf. doi:10.1093/evolinnean/kzag003.

21. Lynch M. Evolution of the mutation rate. Trends in Genetics. 2010;26(8):345–52. doi:10.1016/j.tig.2010.05.003.

22. Sung W, Ackerman MS, Miller SF, Doak TG, Lynch M. Drift-barrier hypothesis and mutation-rate evolution. Proceedings of the National Academy of Sciences. 2012;109(45):18488–92. Available from: https://www.pnas.org/doi/10.1073/pnas.1216223109. doi:10.1073/pnas.1216223109.

23. Mayr E, Ashlock PH. Principles of Systematic Zoology. 2nd ed. New York: McGraw-Hill; 1991.

